# Global patterns of human ageing

**DOI:** 10.1101/124792

**Authors:** Saul J. Newman, Simon Easteal

## Abstract

Ageing is a common, but not universal^1–3^, degradation of biological systems. Ageing in human populations is marked by a highly constrained, log-linear acceleration of the probability of mortality with age^4^. This acceleration determines the onset and duration of human morbidity^5^ and the upper limits of human life. Recent studies have revealed remarkable taxonomic diversity in mortality-derived ageing rates^1,3^. However, the extent of intraspecific variation in ageing rates is assumed to be negligible^2^. Here we show the considerable diversity of human ageing rates, across 81,000 population-specific longitudinal measures of the ageing rate derived from 330 billion life-years of mortality data. These data reveal remarkable flexibility and unexpected trends in the pattern of human ageing. Populations with longer life spans have faster rates of ageing, global ageing rates have doubled historically and women age faster than men. Furthermore, we show that diverse causes of death accelerate at a similar rate with age, that removal of leading causes of death does not alter the ageing rate, and that ageing rates are linked to reproductive schedules in humans. These results challenge accepted ideas in ageing research and provide a broad empirical grounding to study human ageing through population diversity.

The probabilty of death accelerates with age, from the age of sexual maturity, at an approximately log-linear rate in humans and other primates (Fig. 1). This acceleration of mortality is an established indicator of physiological decline and ageing^1,6,7^, expressed as either the time taken for the probability of mortality to double with age (the mortality rate doubling time; MRDT)^6,8^ or the beta exponent of the Gompertz-Makeham^9^ (GM) accelerated failure time model.

**Figure 1.**
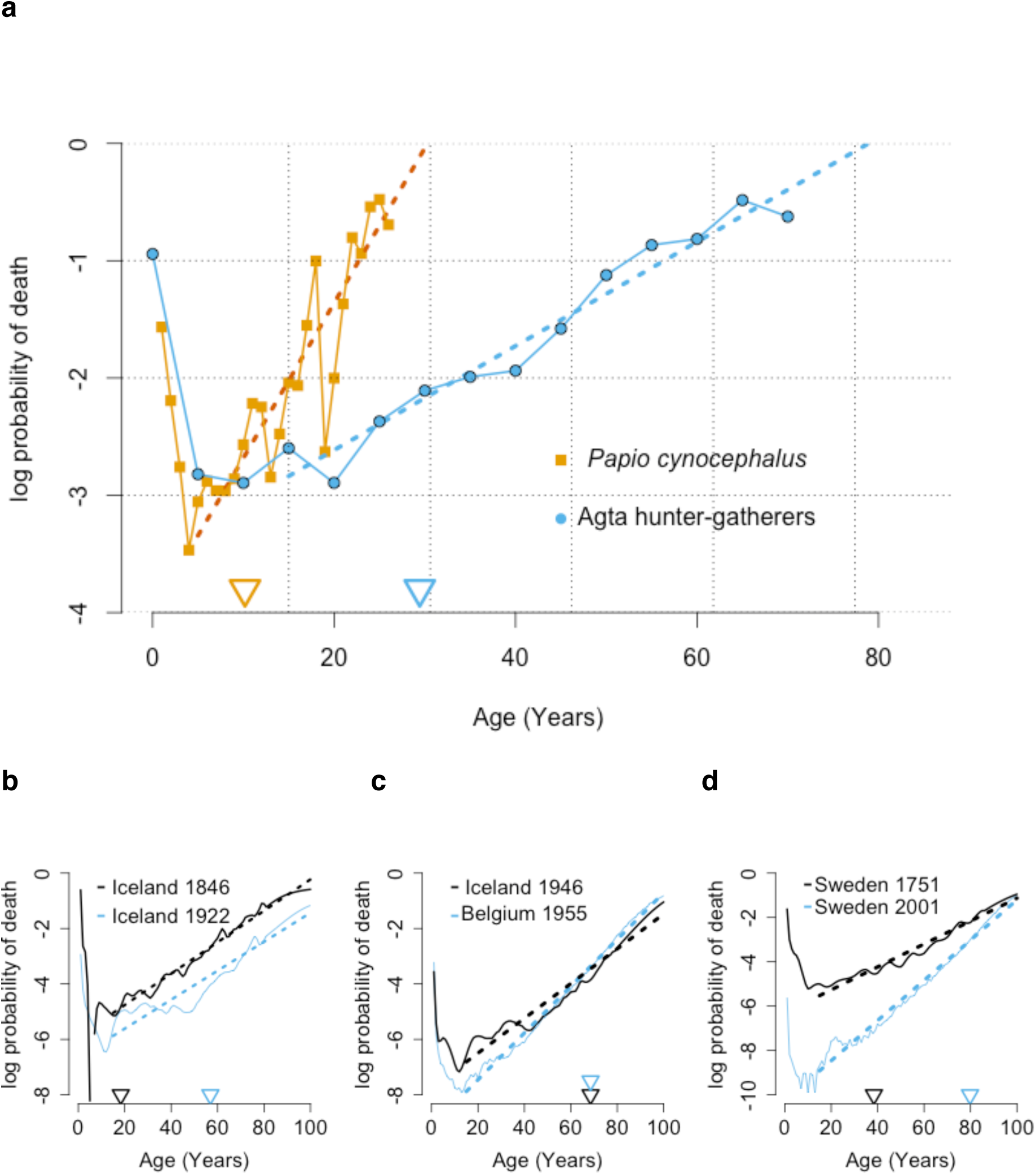
Ageing rate diversity and lifespan. **a**, The ageing rate, indicated by the slope of a robust, log-linear increase (dashed lines) in the probability of death (solid lines). Increased survival and longer average lifespan (triangles) are often thought to result from slower ageing, as shown here for humans (blue) and baboons (orange). However, populations can have: **b** different lifespans but identical ageing rates, **c** identical lifespans but different ageing rates and **d** longitudinal increases in both lifespan and ageing rate.

We calculated MRDT and GM ageing rates using extensive, publicly available life table data from the Human Mortality Database^10^ (HMD), the United Nations (UN) world population prospects database^11^ and Japanese historical data^12^. In addition, we calculated life table data and ageing rates from primate data in Jones *et al*.^1^, Casiguran Agta hunter-gatherer population data^13^, and 39 million deaths located to 3144 United States counties by the Centre for Disease Control (CDC)^14^.

These results highlight considerable variation in human ageing, with clear counterintuitive patterns. For instance, MRDTs exhibit a 4.9-year interquartile range for historical data and a 3.3-year interquartile range across global populations (2010 data). Both models predict observed mortality rates with a high degree of accuracy. However, log-linear MRDT models perform best on average, capturing 95% of variation in mortality rates with half the mean squared error of GM models (SI).

Human populations that live longer, age faster. Longer average lifespans are associated with faster mortality acceleration in all 81,082 population-years tested (MRDT r=0.81; GM rate r = 0.35; p < 2e-16; SI), contemporary national populations (r = 0.84; Fig. 2a), US counties (r = 0.36; Fig. 2b; Extended Data Fig. 1) and historical populations (r = 0.83; Fig. 2c-d).

**Figure 2.**
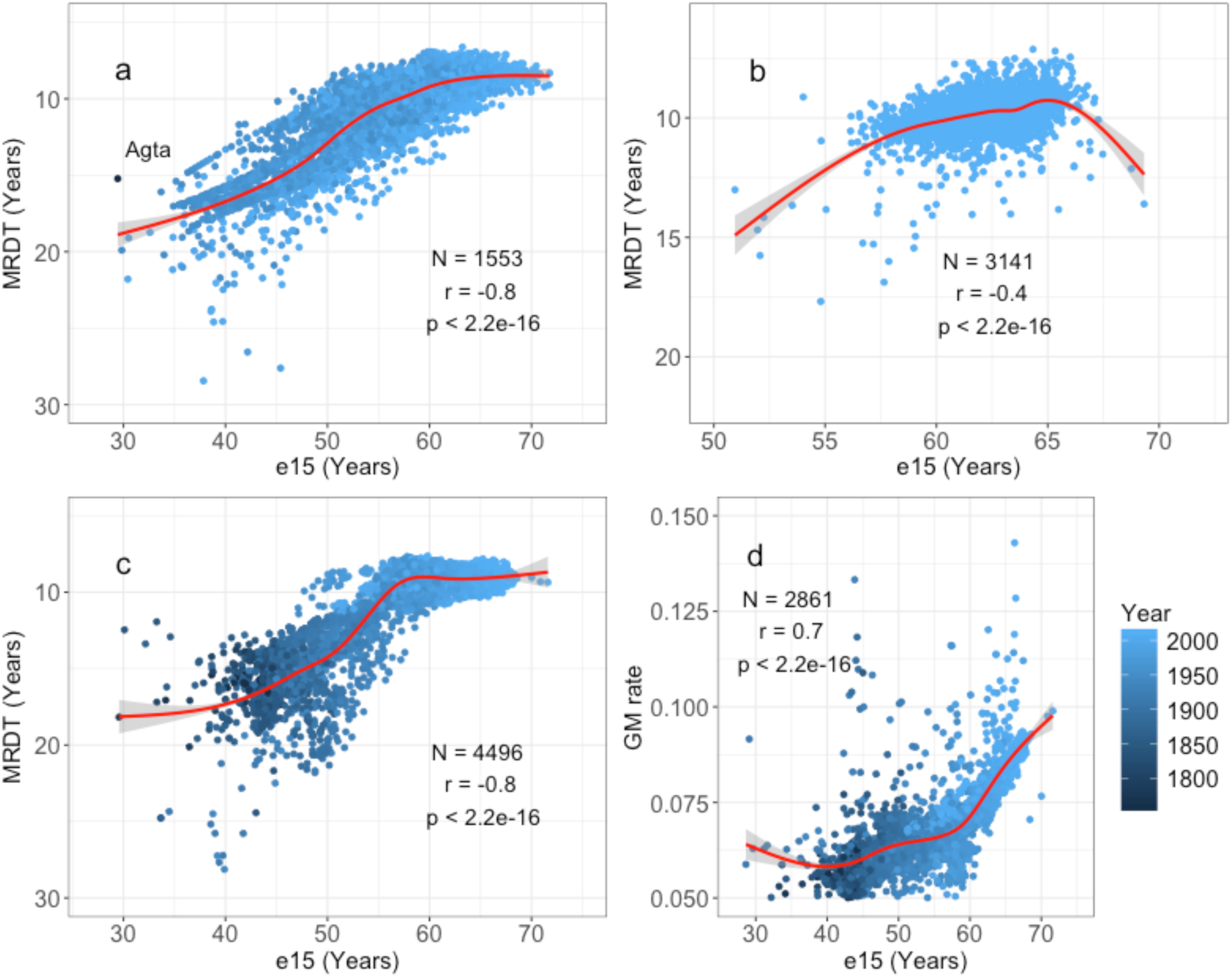
The rate of ageing and adult life expectancy. Positive correlation between life expectancy at age 15 (e15) and the ageing rate, as measured by: **a** MRDT gender-partitioned national populations 1950-2015, **b** MRDT in US counties 1999-2014, **c** MRDT and **d** GM rates in 49 historical populations 1751-2014. Trends fit by bootstrapped locally weighted smoothed splines (red line with 95% CI), showing an inflection point around e15**=**60. Shown in **a** is the Agta Hunter-gatherer population c.1960-2011, excluded are extreme mortality outliers Cambodia c.1975-80 and Iranian males c.1980-85.

The probability of death increases more rapidly from a much lower baseline in long-lived populations. For example, women from Hong Kong have the longest observed average lifespan at 86.5 years, while the Agta hunter-gatherer population average only 25.6 years. Between the ages of 15 and 60 mortality rates increase 29-fold in Hong Kong women and only 6-fold in the Agta.

The relationship between longer lifespan and faster mortality acceleration is highly consistent (Fig. 3a-f). Life expectancy and ageing rates are tightly coupled across 263 years of historical data (Fig. 3a-c), and global increases in lifespan since 1950 are matched by corresponding changes in MRDT and GM rates (Fig. 3d-f).

**Figure 3.**
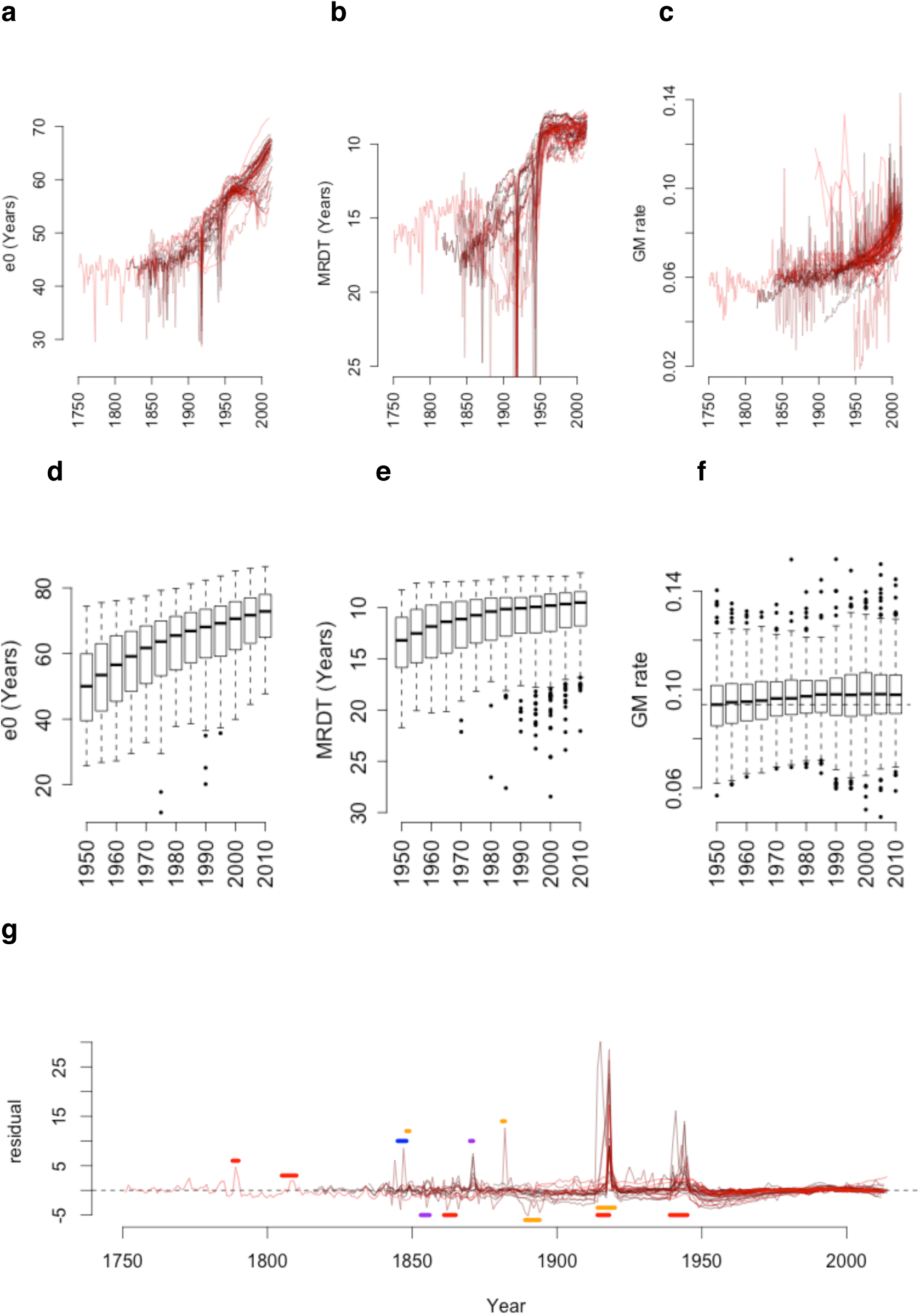
Historical and recent changes in ageing rates. **a-c**, Historical records from 49 populations 1751-2014 and **d-e**, recent records from 205 national populations 1950-2015 show the longitudinal increase of life expectancy at birth (e0), the MRDT and the GM rate of ageing. **g**, Significant deviations from historical MRDT trends are coincident with wars (red), disease outbreaks (orange), concurrent wars and disease outbreaks (purple), and famines (blue) associated with over 100,000 deaths (SI).

Global ageing rates are increasing rapidly, with a 2.9 year faster MRDT and a 12% faster GM rate of ageing since 1950 (Fig. 2, Extended Data Fig. 2). Modern mortality rates double twice as fast with age compared to 19^th^ century rates (Fig. 3b). Ageing rate increases are projected to continue through 2100 (Extended Data Fig. 2). The only significant disruptions of these trends coincide with extreme distortion of early-adult mortality from war, famine and disease events (Fig. 3g).

Shifts in ageing rate do not appear to reflect mortality distortions caused by the removal of early-onset or transmissible causes of death. Rather, ageing rates reflect the parallel increase of the age-specific probability of death across diverse and independent causes (Extended Data Fig. 3). Within populations, almost all causes of death accelerate at a similar log-linear rate regardless of aetiology (Extended Data Fig. 4). For example, the top 15 leading causes of death account for 68% of mortality in the USA, but their collective or individual removal does not significantly change overall ageing rates (SI).

**Figure 4.**
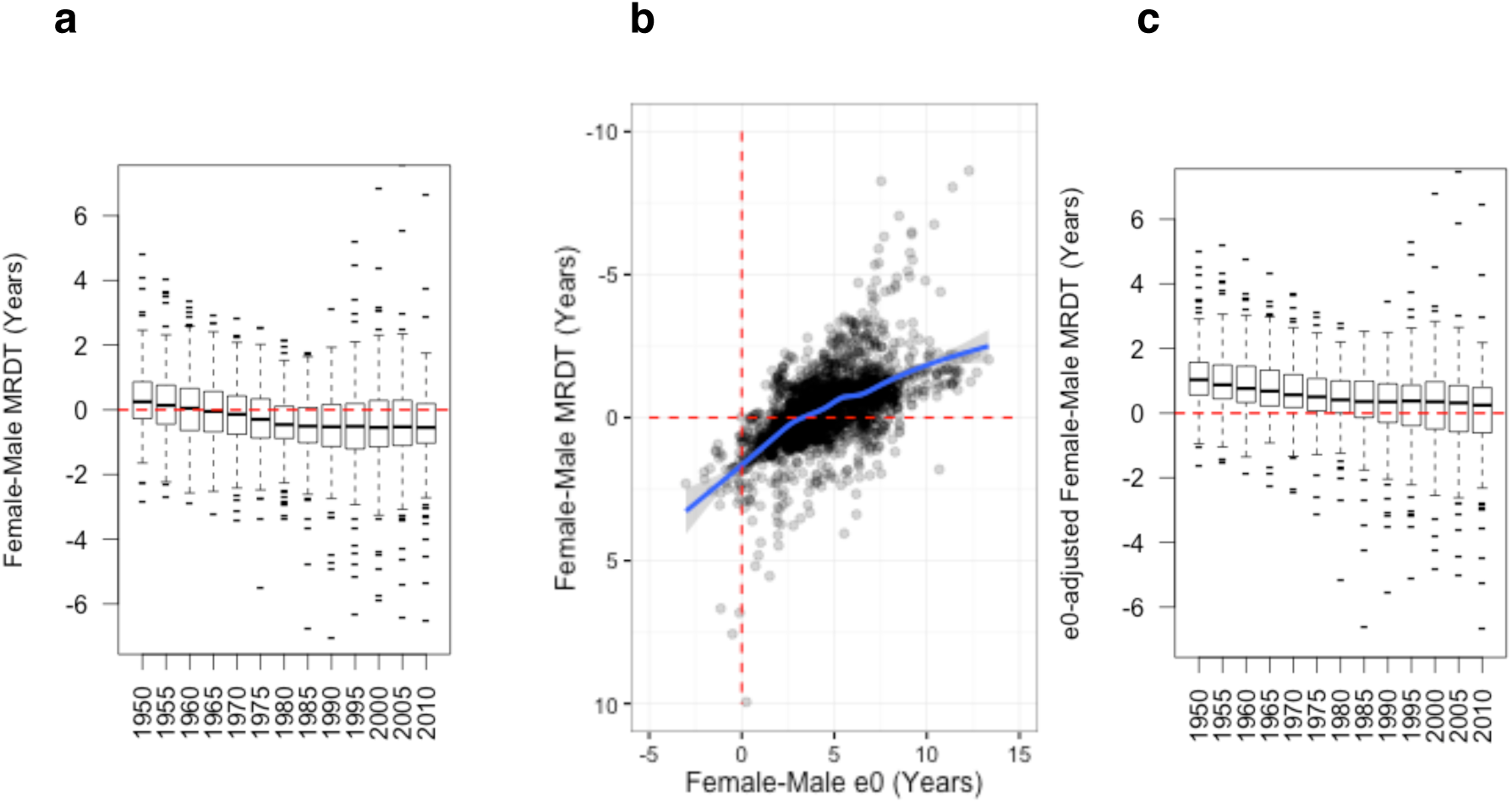
Global sex differences in ageing rates and life expectancy. **a**, Ageing rate differences by sex, showing a transition from faster ageing in men than contemporary women during 1950-1955, to faster ageing in women than men by 2010-2015. **b**, Sex differences in ageing rates and life span, showing the slower ageing rate of women compared to men of an equivalent life span. **c**, Lifespan-adjusted ageing rates by sex since 1950, with a narrowing gap in ageing rates between men and women of equivalent life spans. Red lines indicate gender parity, blue line indicates a locally weighted smoothed spline.

Exceptions to this log-linear pattern of acceleration include extrinsic causes of death such as motor vehicle accidents, and breast cancer (Extended Data Fig. 4). Breast cancer breaks log-linear trends due to a clear sex divide: breast cancer mortality doubles at a typical rate in women (8.8 years) but less than half this rate in men (19.3 years; Extended Data Fig. 4; SI). While the cause of this sex-specific division is unclear, population data reveal systematic differences in ageing rates between sexes.

Women age faster than men. Mortality rates accelerate faster in female populations across 68% of modern nations, with an average 0.6-year faster MRDT in women than men of the same nation (p = 3e10^−16^; Fig. 4a; SI). A large proportion of this gender gap is explained by the 4.5-year shorter average male lifespan (r = 0.6; Fig. 4b). However, adjusting for lifespan does not fully resolve sex differences in ageing rate.

In a majority of populations women age more slowly than men of an equivalent lifespan. However, this imbalance is narrowing globally (Fig. 4c). Women age faster than men of equivalent lifespan in high-income countries, and by 2030 women are projected to have both faster lifespan-adjusted and overall ageing rates than men worldwide (Extended Data Fig. 2).

Selection for later ages at reproduction leads to slower ageing in laboratory populations^15^, suggesting some ageing rate diversity may be explained by the relative timing of reproduction. Faster absolute and lifespan-normalized ageing rates are linked to lower total fertility rates, earlier reproduction and slower declines in fertility rate with age in national populations (Extended Data Fig. 5a-b). In contrast to model organisms^15^, later reproduction is associated with faster ageing. This trend held across US counties, with a later onset of reproduction associated with more rapid ageing for absolute (r = 0.4; p<2.2e^-16^; Extended Data Fig. 6a-b) and lifespan-normalized ageing rates (r = 0.4; p<2.2e^-16^; Extended Data Fig. 6c-d).

We considered the hypothesis that mortality accelerations are independent of physiological rates of ageing, but found this hypothesis difficult to assess. While death is an easily scored phenotype available for billions of individuals, no equivalent biomarker of ageing exists. However, 27 CDC indicators of healthy ageing from 51 US states and territories^16^ support the use of mortality indicators of ageing. Ageing rates from 20 CDC indicators show faster ageing in long-lived populations and 11 indicators are positively correlated with MRDT (corrected p-value<0.05; SI).

These results depart from findings of previous ageing rate studies. In particular, the positive correlation between lifespan and ageing rate suggests the need to reconsider proximate causes of ageing. This discrepency appears to reflect misunderstanding of the statistical differences between survival curves, lifespan and ageing. It is often assumed incorrectly that longer lifespan or higher survival rates indicate slower ageing (*e.g*. Kuro-o *et al.*^17^; Kenyon^18^). However, factors that modify survival and lifespan do not necessarily affect ageing rates.

Increased lifespan and survival rates may reflect slower ageing^19^, reduced infant or child mortality rates^6^, the delayed onset of ageing^19^, lower extrinsic mortality^19^ (Fig. 1b) or any combination of these factors (SI). Comparison of the average or maximum lifespan, and survival analyses such as Kaplan-Meier test statistics, have no power to discriminate between these effects.

Detection of differential ageing rates usually requires a sample of at least 1000 deaths per population ^20,21^, rising to 10-20 thousand deaths for late-life changes^21^. Ageing rates can be measured in smaller populations if strict assumptions are met^4^, but generally suffer serious bias and inflated variance below this threshold (Extended Data Fig. 7).

As a result studies that measure ageing rates, as distinct from longevity or survival, can reveal novel patterns. For instance, humans age at similar or faster rates than shorter-lived primates^6,22^ and outlive species with slower or negative rates of ageing^1,3,23^.

Such studies have revealed the absence of mortality accelerations in other species^1,3^, clearly indicating that ageing responds to, and can be reduced by evolutionary pressure. This study further reveals considerable flexibility in ageing rates across short timescales and diverse environments, suggesting the potential to environmentally modify ageing rates within bounds set by selective landscapes.

A moderate one-year reduction of global MRDT, well within the current range of human variation, would be sufficient to increase average lifespan by 10 years (SI). Understanding the proximate and ultimate drivers of mortality acceleration, and the cause of their high level of diversity in humans, is therefore of considerable interest for both ageing research and medicine.

## Methods

### Life table construction

Historical mortality data and life tables were downloaded from the Human Mortality Database^10^ (accessed 28 April 2016). Historical Japanese male-female segregated population data was obtained from life tables supplied in the ‘fmsb’ package^12^ version 0.5.2 in R^24^. National life table data were obtained from the United Nations (UN) world populations prospects^11^ for the period 1950-2010 (observed) and 2010-2100 (projected).

United States county data on mortality rates, population size data and cause of death data were downloaded from the Centre for Disease Control (CDC) Compressed Mortality Files, using the CDC Data Access Portal^14^ accessed 5 July 2016 and 15 April 2016. Compressed mortality file data represent 39 million deaths in the United States and territories occurring between 1968 and 2014, partitioned by sex, race, age, cause (before 1999) and location of death.

Data was excluded for the year 1972, which constituted a 50% sample of all deaths^14^. Reduced sample sizes during this year were associated with transient shifts in life expectancy and ageing rates, consistent with ageing rate biases introduced by small population sizes (Extended Data Fig. 7).

Standard life tables were calculated for US county data using the fmsb package^12^ version 0.5.2 in R^24^, on the basis of supplied age-specific probabilities of death (*q_x_*) estimates. Deaths occurring before one year of age were pooled to standardize the infant mortality rate, despite the availability of higher resolution neonatal mortality data. All life table data were calculated to account for the partial non-standard annual, quinquennial and decennial pooling of age categories within US county data^14^.

As age-specific mortality rates (*q_x_*) were not reported in the US county data from 1968-1998, we calculated these values from reported ages at death and the population estimate of total residents reported for each county^14^. Deaths were assumed to be symmetrically distributed within age categories, in line with US county data from other years^14^ and best practice^10^.

In US county data from 1968-1998, the probability of death within each age group is ascertained during a single year. However, age categories are variously pooled decennially, quinquennially, and annually^14^. We therefore approximated the probability of survival across age categories by multiplying the observed probability of death during a single year by the number of years of each age category.

Rather than being redistributed into age categories, individuals with uncertain ages at death in 1968-88 US county data (0.03%; N=13367 deaths) were excluded from analysis. This practice aligns with omission of uncertified deaths in national reporting for the UN data^11^ and the World Health Organization^25^.

The exclusion of deaths of unknown age had minimal impact on mortality estimates. On average only 5.4 deaths of uncertain age, out of an average 13,000 deaths, were recorded per county-year. This incurred a maximum underestimate of 6.5% (3 of 54 deaths missing) and an average 0.04% underestimate in the age-specific mortality rate for any age category.

Life table data for the Agta population was calculated from the observed age-specific probability of mortality calculated by Newman & Easteal^26^ using the ‘fmsb’ package^12^ in R^24^ version 3.2.1. Data on the genealogies, birth dates and death dates in this population are available for direct download from the Agta Demographic Database^13^, and the calculated life table is available on request.

Individuals with unknown years of death were excluded, and we assumed deaths were equally distributed within age categories. In contrast to work on other hunter-gatherer populations, no models were fit to approximate the age-specific probability of death for these data.

### Calculation of ageing rates

The rate of ageing was calculated from collected life table data for all unique population-years, using two widely accepted ageing models fit to the log-transformed age-specific probability of death.

Ages where the observed probability of death was zero, or where the probability of death is forced to equal one in terminal age categories^10^, were excluded from model fitting. Populations were excluded from analysis if they were missing more than 10% of *q_x_* data, or in the case of US county data, where more than one data point was missing.

The mortality rate doubling time (MRDT) was estimated by fitting an ordinary least squares linear model to log(*q_x_*) data, and calculated as 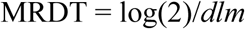; where *dlm* is the slope of the linear regression. The Gompertz-Makeham rate of ageing (GM rate) was estimated by fitting a Gompertz-Makeham mortality model^4,9^ to quinquennial log(*q_x_*) values using the fmsb package^12^ version 0.5.2 in R^24^, returning the beta exponent using default initialization parameters.

Within the HMD, all period data have been fit by smoothed cubic splines to approximate population size^27^ and by the Kannisto old-age mortality model^27^ above the age of 85. Likewise, a large proportion of US mortality data are pooled above this age^14^ into a terminal “85+” or “80+” age category. Therefore, we fit ageing models across ages 15 to 75 years inclusive, to avoid model-approximated data and to allow more direct comparison of ageing rates across datasets.

Gender differences in ageing rate were measured using MRDT and GM rates for individuals of the same national population and year in the UN national data^11^ and the Japanese historical^12^ dataset.

The relationship between gender-pooled ageing rates and adult life expectancy from age 15 years (e15) was fit by a locally weighted smoothed spline^28^ in R^24^ (Fig. 4b). This smoothed model was used to predict the expected rate of ageing for the given adult life expectancy within each gender-discriminated national population.

The age of onset of ageing was estimated by fitting a bootstrapped nonparametric nonlinear regression between the log probability of death *q_x_* and age using the *np* package^29^, and predicting the minimum age-specific probability of death (Extended Data Fig. 8). All ageing rates are included in the supplementary information (SI).

### Partitioning by cause of death

United States county data from 1968-2014 is certified according to the international cause of death certificates (ICD-9 and ICD-10) and recoded to a longitudinally consistent set of 69 (1968-1978), 72 (1979-1988) and 113 (1989-2014) cause-of-death meta-categories^14^.

Data is available at the county level prior to 1989 for ICD-9 (1968-1979) and ICD-10 (1980-88) raw cause-of-death codes, re-coded to 72 longitudinally comparable causes of death. We measured the age-specific rate of mortality within each of these codes for compressed mortality data, for populations aggregated by race and sex, and pooled these results by state or US territory. The probability of death at each age was approximated from the reported population size within each quinquennial or decennial age category.

The probability of mortality was log-transformed and fit by least squares regression across ages 15-75 years for each cause of death, and the rate of mortality doubling with age calculated (Extended Data Fig. 4). State-year combinations where less than a thousand deaths occurred were screened from these samples, as this sample size was insufficient to accurately calculate MRDTs (Extended Data Fig. 7).

The effect of removal of causes of death was simulated by individually and collectively removing the 15 leading causes of death during 1988 from mortality counts, and recalculating MRDT from the remaining deaths across each US county (SI). There were a minimum 700 counties with valid recalculated MRDTs for each mortality indicator after this process. The distribution of MRDT values was compared between the baseline and modified rates using a student’s t-test.

### Environmental correlates

Quinquennially pooled data on age-specific fertility rate, gross reproductive rate and total fertility rate were downloaded from the World Populations Prospects^11^ for all national populations since 1950. County-specific fertility indicators were downloaded from the US census bureau^30^. The early to late fertility ratio (Extended Data Fig. 5b) was calculated as the ratio of age-specific fertility rates before and after age 30.

Indicators of healthy ageing were downloaded from the CDC^16^. Rates of ageing were calculated for CDC healthy ageing indicators using the same general approach as the MRDT. Age-specific rates were log-transformed, fit by a linear least squares model, and reduced to a doubling time (for negative indicators of health, *e.g*. disability rates) or half-life (for positive indicators of health, *e.g*. ‘good or very good’ self-reported health) to estimate the rate of physiological ageing.

We then measured the correlation between these physiological ageing rates and the state or territory-level rate of mortality-derived ageing, measured by MRDT and GM rates. Correlation coefficients were Bonferroni corrected by the total number of comparisons.

## Data Availability

The authors declare that all data are available within the paper and its supplementary files.

## Author contributions

S.J.N. wrote the analysis and code, extracted and quality-checked the data, and performed the initial analysis. S.J.N. and S.E. developed the study concept, analysis and statistical design, performed the analysis and co-wrote the manuscript.

---

**Extended Data Figure 1.**
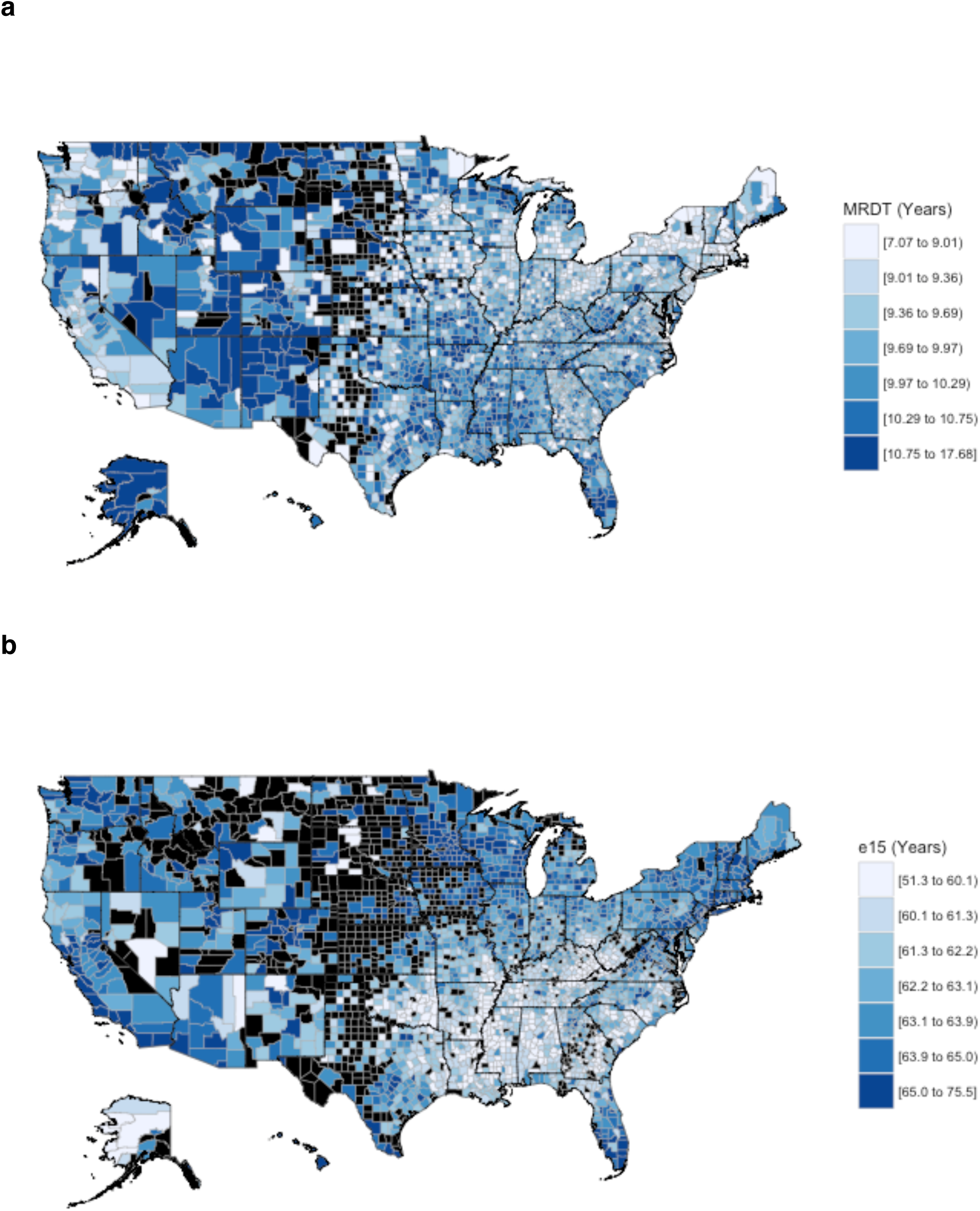
Adult life expectancy and ageing rates across US counties 1999-2014. **a**, The rate of mortality doubling and **b**, adult life expectancy from age 15 years (e15), measured across 3141 US counties from 1999-2012. Darker blue indicates **a**, slower rates of ageing and **b**, longer life span. Blacked-out counties contain insufficient data (missing data or N<1000 deaths) to accurately calculate ageing rate or life expectancy.

**Extended Data Figure 2.**
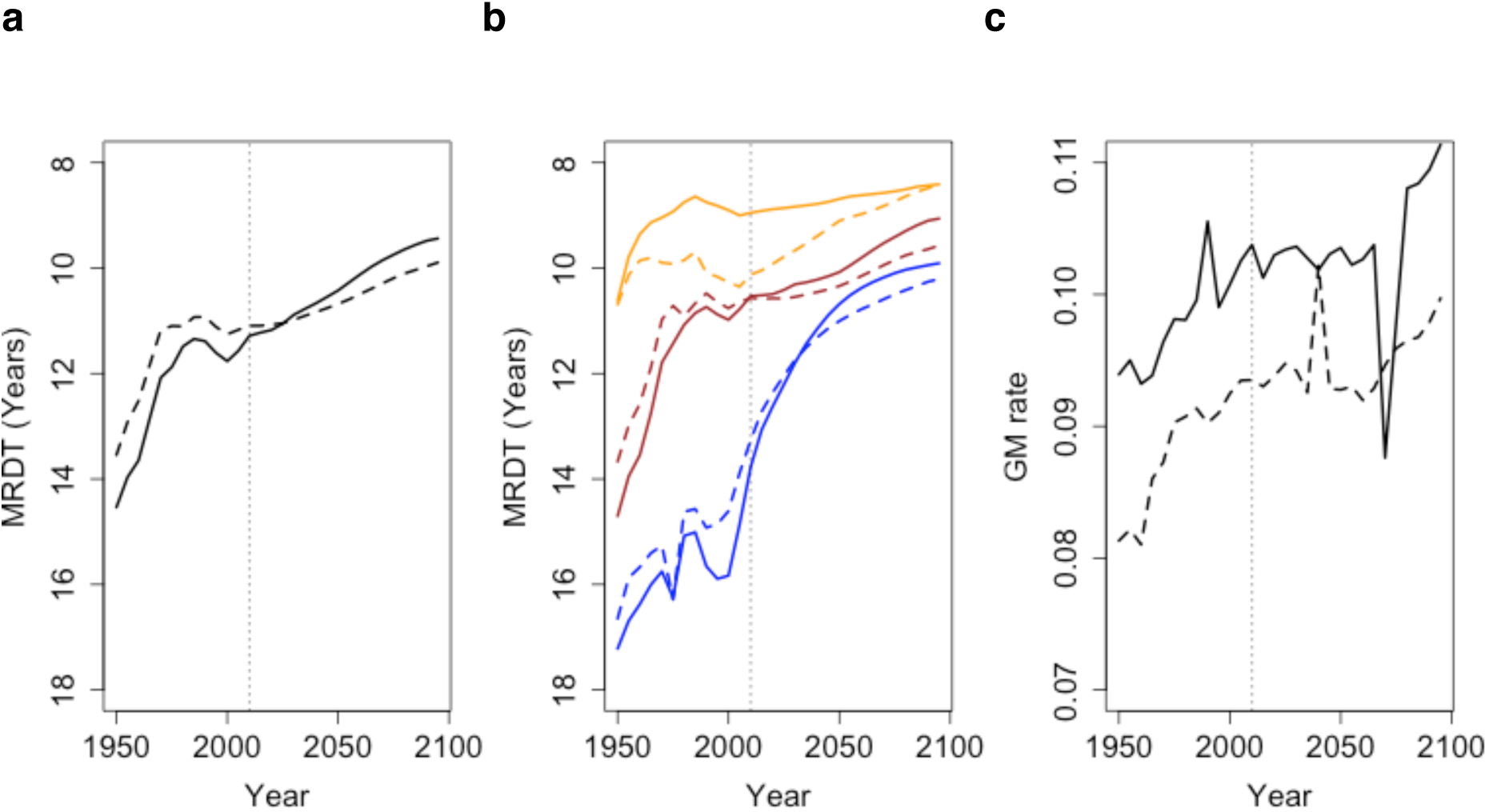
Observed and projected changes in ageing rates 1950-2100. **a**, Global and **b**, income-partitioned trends in MRDT indicate ongoing increases in the rate of ageing to 2100 for both women (solid lines) and men (dotted lines). **c**, Similar trends occur in the Gompertz-Makeham rate of ageing. Dotted vertical lines separate projected and observed data; orange, red and blue lines represent pooled UN high, medium and low-income countries respectively.

**Extended Data Figure 3.**
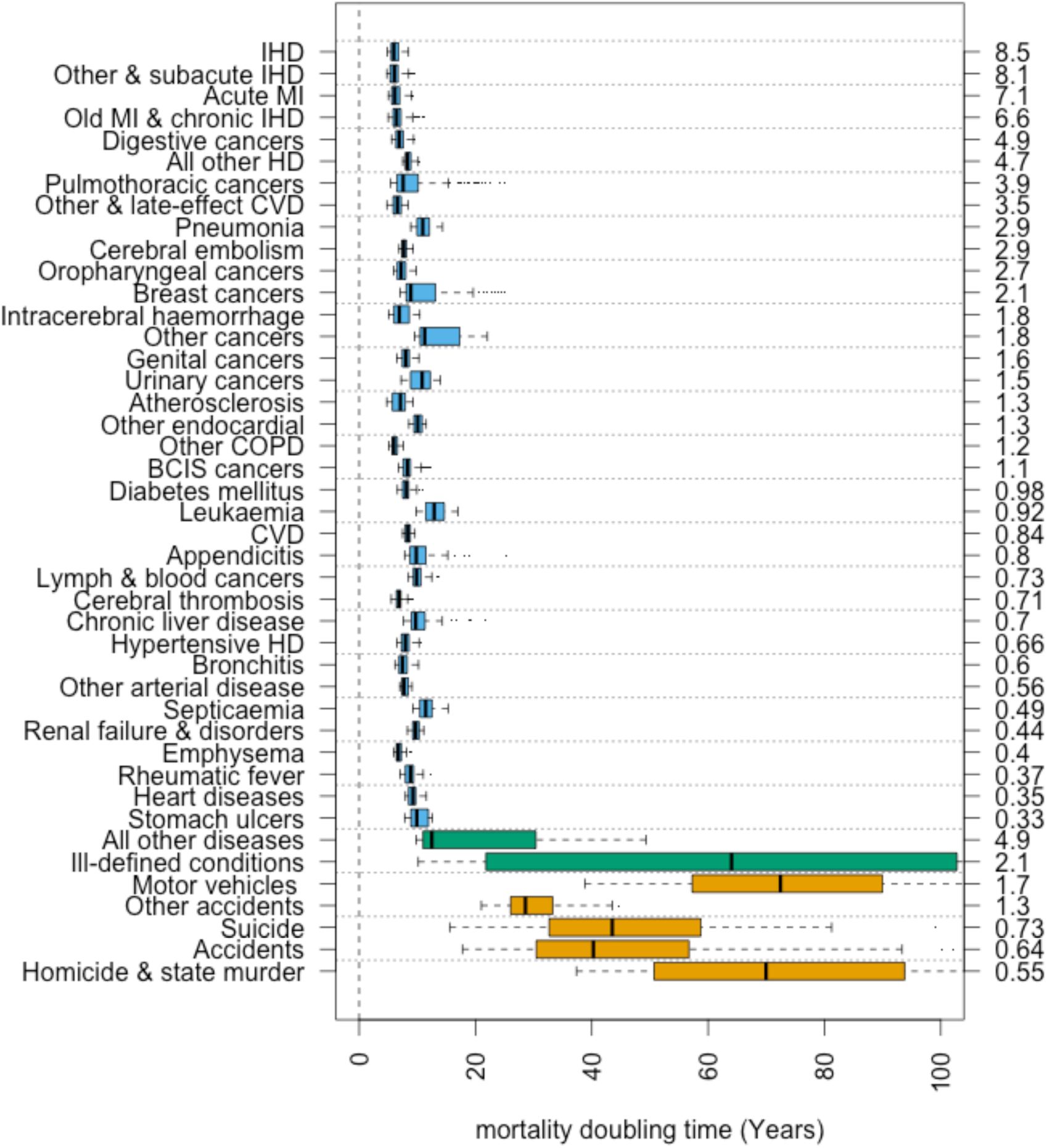
Acceleration of leading causes of death with age. The doubling time of mortality rates with age from 43 CDC-coded causes of death, measured across 1132 US state-years from 1968-1988. Despite a long census period, the probability of mortality doubles every 6-10 years of age within most categories, independent of proximate cause or disease aetiology. Extrinsic causes of death (orange) and poorly defined categories (green) accelerate much slower, and have poor fit under log-linear models. Y-axes indicate the cause (left) and percentage (right) of deaths. Acronyms are ischemic heart disease (IHD), heart diseases (HD), cerebrovascular disease (CVD), chronic obstructive pulmonary disease (COPD) and benign carcinomas, carcinoma in situ and neoplasms of uncertain behavior and unspecified nature (BCIS).

**Extended Data Figure 4.**
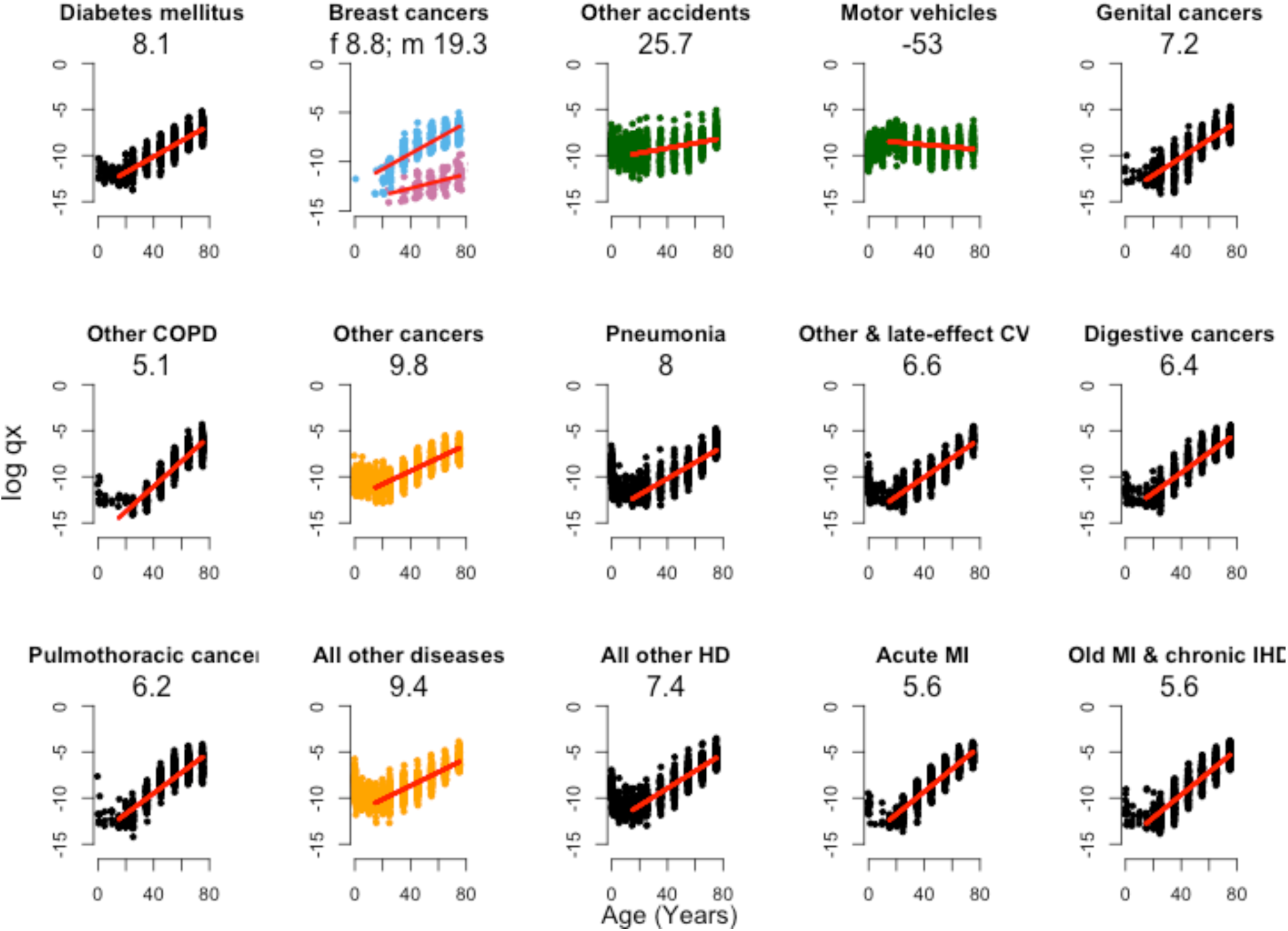
Acceleration of the top 15 causes of death in 1988 for the USA. The probability of death from the 15 leading causes of mortality in US adults, measured for counties with over 1000 deaths. Numbers indicate the doubling time (positive) or half-life (negative) of the probability of death with age, in years. Log-linear increases in the probability of death are apparent beyond age 15 for all categories except breast cancer, which is partitioned by male (pink) and female (blue) rates, and extrinsic causes of death (green). Acronyms as in Extended Data Fig. 3.

**Extended Data Figure 5.**
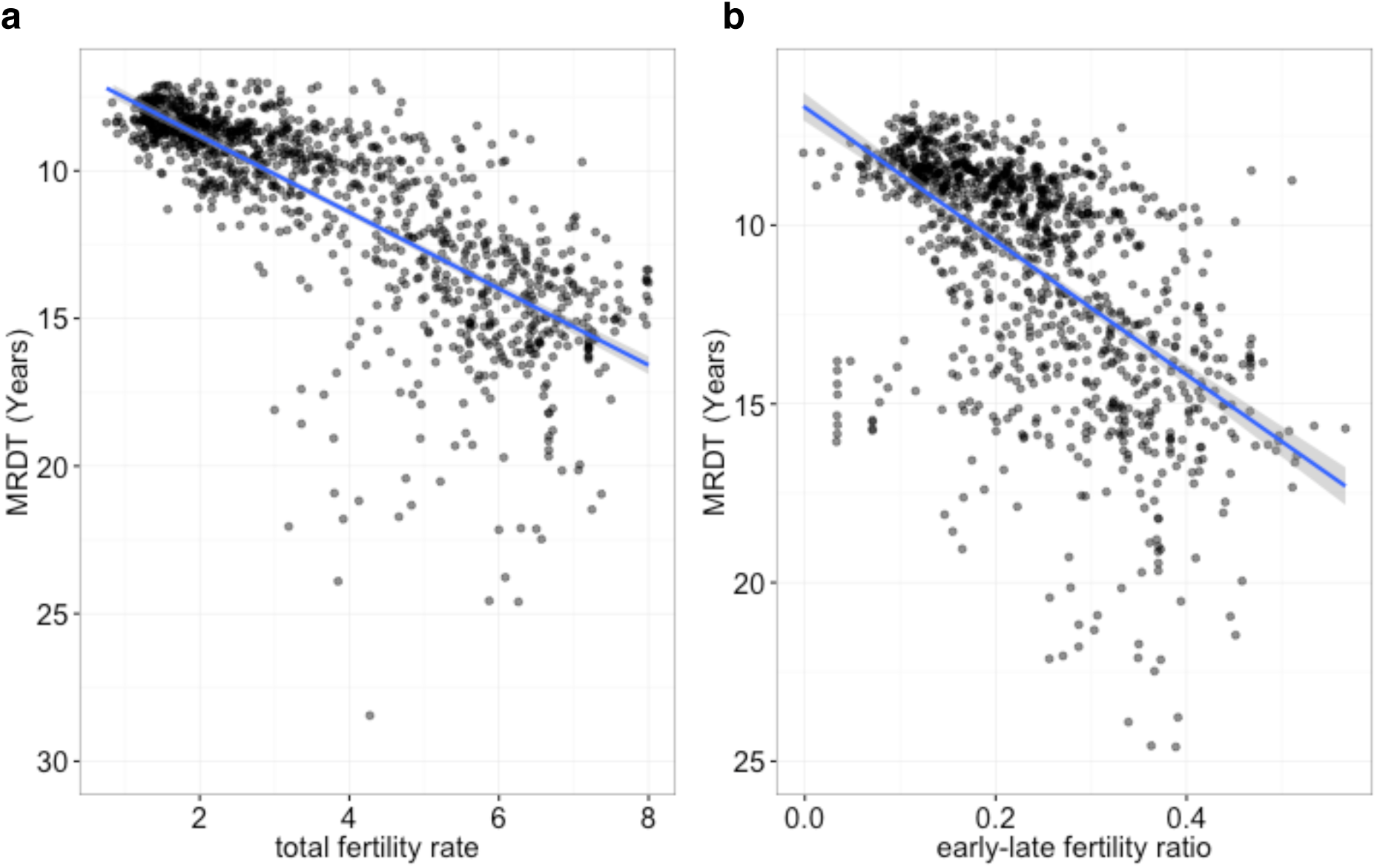
Ageing and fertility rates in national populations. **a**, In UN national data, higher total fertility rate is correlated with slower ageing rates in both men (r = 0.66) and women (r = 0.73; shown). **b**, A similar trend occurs for the ratio of fertility rates before and after age 30, with higher relative fertility at older ages linked to faster ageing (r = 0.54).

**Extended Data Figure 6.**
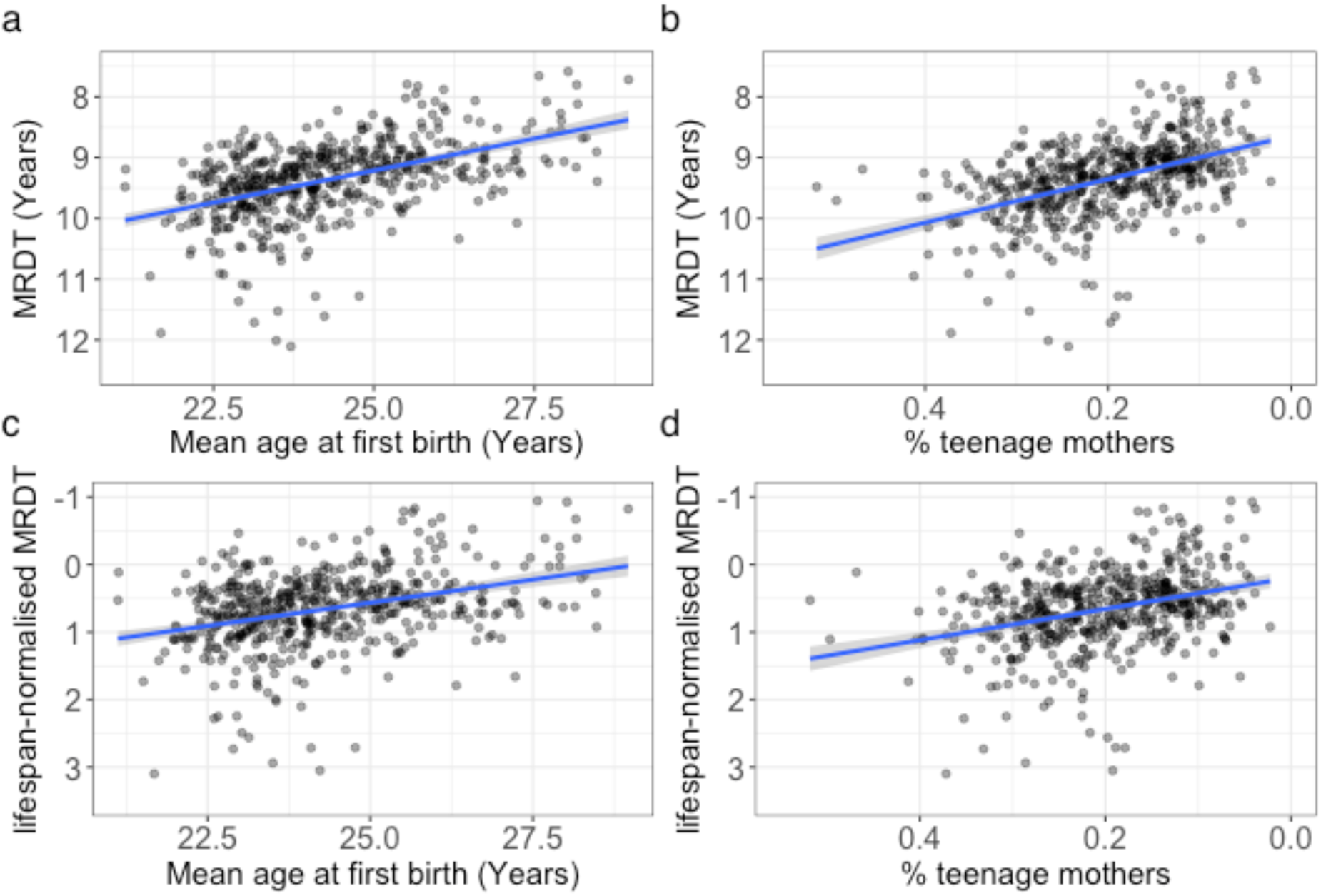
Ageing rates and the initiation of reproduction. Across 573 gender-pooled county populations with complete mortality and reproductive data, faster ageing rates were associated with **a**, a later mean age at first birth (r = 0.36; p <2.2e-16; N=521) and **b**, lower teenage birth rates (r = 0.55; p <2.2e-16; N=521), with lifespan-adjusted ageing rates showing similar trends for both **c**, earlier reproduction (r = -0.4; p <2.2e-16; N=513) and **d**, higher teenage birth rates (r = -0.4; p <2.2e-16; N=514).

**Extended Data Figure 7.**
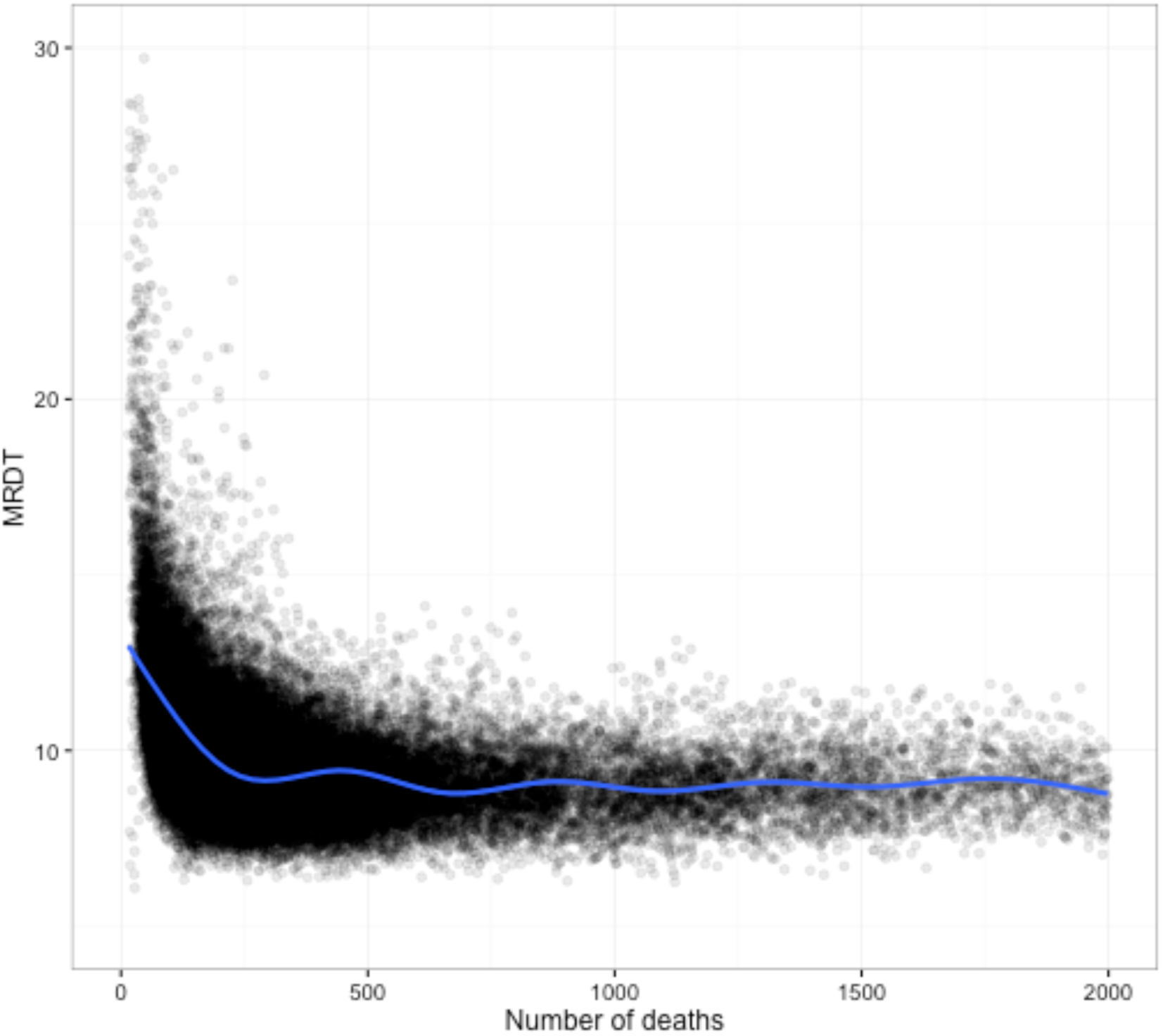
Population size biases in ageing rate. Ageing rate measurements, here shown by mortality rate doubling time, suffer a strong positive bias and increasing variance when measured in populations with a small number (<1000) of observed deaths.

**Extended Data Figure 8.**
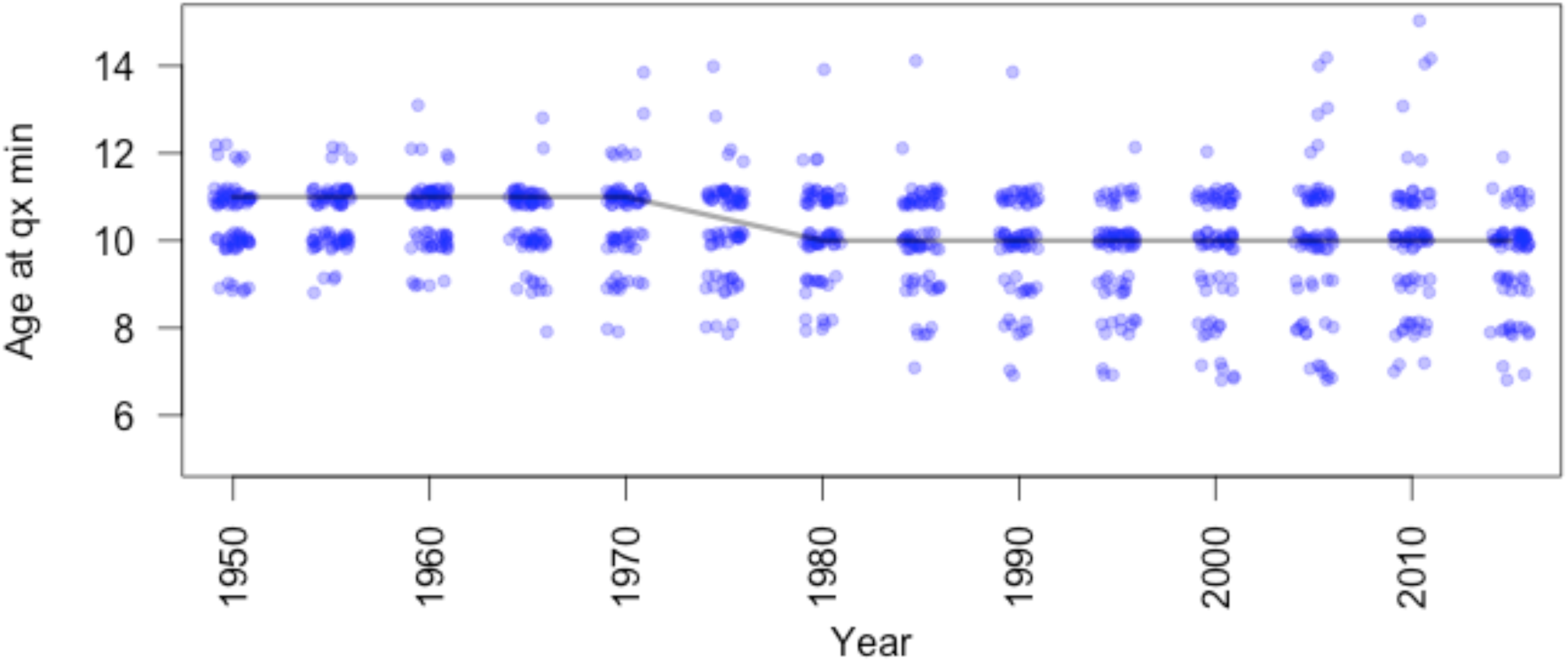
Change in the median onset of ageing since 1950 in UN data. Data show a one-year reduction in the onset of ageing since 1950 (qx min; black line). However, this pattern may reflect increased sampling noise in the probability of mortality at these ages, caused by considerable falls in the probability of death across this census period.

